# Spatial Transcriptomics Using Archived Formalin-Fixed Paraffin-Embedded Core Needle Biopsy Tissues Revealed Unique Transcriptomic Signatures in Kidney Transplant Rejections

**DOI:** 10.1101/2025.04.08.647600

**Authors:** Yan Chen Wongworawat, Chirag Nepal, Mark Duhon, Wanqiu Chen, Minh-Tri Nguyen, Adam Godzik, Xinru Qiu, Wei Vivian Li, Gary Yu, Rafael Villicana, Craig Zuppan, Michael De Vera, Mark Haas, Charles Wang

## Abstract

Clinical core needle biopsies present challenges for transcriptomic analysis due to limited tissue volume. In this study, we determined the feasibility of using spatial transcriptomics to evaluate rejection on formalin-fixed paraffin-embedded (FFPE) core needle biopsies from human kidney allografts. We demonstrated that non-rejection, active antibody mediated rejection (AMR), acute cell mediated rejection (TCMR) and chronic active AMR have distinct transcriptomic signatures. Subclusters of monocytes/macrophages with high *Fc gamma receptor IIIA (FCGR3A)* expression were identified in C4d-positive active AMR and acute TCMR, and the spatial distribution of these cells corresponded to the characteristic histopathological features. Key markers related to monocyte/macrophage activation and innate alloantigen recognition were upregulated, along with metabolic pathways associated with trained immunity in AMR and TCMR. The discovery of unique transcriptomic signatures associated with AMR and TCMR facilitates the differentiation of acute kidney allograft rejections using spatial transcriptomic data derived from FFPE core needle biopsies.

## 1. Introduction

Allograft biopsy remains the gold standard for diagnosing kidney transplant rejections. International standard classification systems, Banff classification, define a set of histopathologic features that indicate the type and severity of rejection in post-transplant biopsies. However, the empirical histology system is not always accurate and may miss subtle or early signs of rejection, misclassify rejection subtypes, or produce false positive results. Transcriptome microarrays such as Molecular Microscope (MMDX) ^1^ and the Banff Human Organ Transplant Gene (B-HOT) panel ^2^ overcome some of the weaknesses of histology via specific transcripts that are associated with different rejection types. Hence, the 2013 Banff meeting adopted a molecular diagnostic option into the classification ^3^. However, MMDX requires additional fresh biopsy cores, limiting its clinical utility of studying archived specimens to track disease progression at the molecular level ^1,2^. There is a significant possibility that the extra core obtained for molecular profiling may not correspond to the original samples examined for histopathological features, which can cause discrepancy ^4,5^. B-HOT can be utilized with formalin-fixed paraffin-embedded (FFPE) tissue samples using the NanoString nCounter platform. However, obtaining sufficient tissue volume for these bulk transcriptome microarrays can be challenging when using clinical core needle biopsies. Additionally, these methods have the limitation of losing anatomic localization or spatial context. Research implementing single-cell RNA sequencing (scRNA-seq) in transplantation studies offers valuable insights into cellular heterogeneity at single-cell resolution. However, this technique has limitations: it typically utilizes fresh or frozen tissues, requires a high number of isolated cells that are difficult to obtain through clinical core needle biopsies, and results in the loss of spatial information ^6^. Spatial transcriptomics can overcome this limitation, by detecting RNA expression and mapping gene activity within a single hematoxylin and eosin-stained (H&E) - stained section from FFPE tissue while preserving spatial context, revealing the distribution of various cell types and molecular pathways within their native microenvironments. This spatial information is particularly valuable in complex tissues like kidney allografts, where the location of immune cells relative to specific kidney structures can provide important diagnostic insights. However, there is a paucity of research implementing spatial transcriptomics in transplantation studies, with the few existing articles exclusively utilizing fresh frozen tissues except one publication studying two cases of TCMR ^7–9^. The morphological preservation of frozen tissue is suboptimal for histopathological examination and less than ideal for spatial transcriptomic colocalization studies. The spatial transcriptomic signatures of FFPE core needle biopsy tissue from human kidney allografts have not yet been fully characterized. Furthermore, it remains to be investigated whether the molecular insights provided by spatial transcriptomics can more effectively differentiate and diagnose between various subsets of rejection and distinguish rejection from other causes of renal dysfunction, such as acute tubular injury (ATI), acute calcineurin inhibitor (CNI) toxicity, and infection.

Increasing evidence highlights the essential roles of innate immune cells, especially monocytes/macrophages and natural killer (NK) cells, in solid organ transplantation ^10–16^. Macrophages play pivotal roles in the innate immune response to transplant allografts during acute rejection by producing proinflammatory cytokines and generating reactive oxygen and nitrogen species (ROS and RNS) ^12,17^. Both donor- and recipient-derived monocytes/macrophages activate adaptive immune responses by functioning as antigen-presenting cells (APC). They activate T cells through co-stimulatory signals, leading to release of pro-inflammatory cytokines and resulting in acute rejection ^15^. Macrophages are also implicated in chronic rejection and graft failure ^15,18,19^. However, the molecular and spatial characteristics of monocytes/macrophages in kidney allograft rejection have not yet been investigated using spatial transcriptomics on FFPE tissues.

In this study, we follow the hypothesis that antibody mediated rejection (AMR) and cell mediated rejection (TCMR) of kidney allograft will demonstrate unique transcriptomic signatures that could be recognized in the spatial transcriptomic data. To test our hypothesis, we conducted spatial transcriptomics on FFPE core needle biopsy samples from human kidney allografts representing various rejection groups to identify distinct monocytes/macrophages subclusters. Additionally, we performed functional pathway and gene network analyses to elucidate the underlying biological, cellular, and molecular processes, especially of innate immunity.

## 2. Materials and methods

### 2.1 Perform spatial transcriptomics FFPE core needle biopsies of human kidney allografts

We performed 10x Genomic Visium spatial transcriptomics analysis on H&E - stained sections from archived FFPE core needle biopsies of human kidney allografts following Visium Spatial Gene Expression for FFPE workflow (Graphic Abstract). 1) Sample preparation and RNA quality control: section FFPE tissues onto charged glass slides. 2) Assess RNA integrity using methods Distribution Value 200 (DV200): DV200 represents the percentage of RNA fragments that are longer than 200 nucleotides in a sample. This method is particularly useful for evaluating the quality of degraded RNA samples, such as those extracted from FFPE tissue. Only samples with a DV200 value equal to or greater than 30% were processed. 3) Performed standard H&E staining directly on the glass slides. 4) Evaluated H&E staining slides to select areas of interest for 6.5 x 6.5 mm capture areas. 5) Probe Hybridization with whole transcriptome probe panels. 6) Used the Visium CytAssist instrument to precisely transfer bound probes onto the Visium slide. The Visium slide contains 6.5 x 6.5 mm capture areas with 55 μm barcoded squares. 7) Generated gene expression libraries from each tissue section (library preparation). 8) Sequenced the libraries on compatible Illumina sequencers, such as NovaSeq 6000 systems. 9) Employed Space Ranger software for data processing, applied standard quality control metrics to filter out low-quality spots, and combined all eight samples into a unified dataset ^20,21,22^. 10) Utilized Loupe Browser for interactive data exploration, integrating whole transcriptome analysis with precise spatial information from archived FFPE samples.

### 2.2 Differential gene expression, clusters identification and cell typing

All bioinformatics analysis was performed utilizing the BioTuring Lens platform (https://bioturing.com) ^23,24^. 10X Visium spots were clustered via the Louvain method (principal component analysis (PCA) Resolution=1). Uniform Manifold Approximation and Projection (UMAP) visualization or t-distributed stochastic neighbor embedding (t-SNE) dimension reduction were generated via PCA of gene expression with no batch correction (n_neighbors=30). Cell types and subtypes per Louvain-derived cluster were predicted using the HaiTam algorithm (https://talk2data.bioturing.com/predict). Spots that were not confidently characterized into a single cell type (i.e., undefined) were omitted from the analysis. Differential expression of genes (DEG) among spots in each case was calculated via the Venice algorithm (*p*<0.05) treating each spot as an individual sample data point. Hierarchical clustering heatmaps of the DEGs were generated and organized via a dendrogram of the cases and cluster plots of marker genes per cluster. Expression of specific genes per spot was measured and overlayed onto the UMAP or t-SNE.

### 2.3 Assessing concordance between FFPE tissue transcriptomic signatures and published RNA signatures of transplant rejection

To evaluate the consistency between our findings and existing research, we compared the transcriptomic signatures of AMR and TCMR from our FFPE tissue analysis with RNA signatures derived from frozen tissue bulk transcriptome microarrays, as reported by Halloran *et al*. in 2017 and 2024. This comparison was visualized using a Venn diagram, highlighting similarities and differences between the two approaches.

### 2.4 Functional pathway and gene network analysis

To analyze the gene networks, canonical, and bio-functional pathways, we applied Gene Ontology (GO) Enrichment Analysis tools to the lists of differentially expressed genes (ShinyGo v0.66, http://bioinformatics.sdstate.edu/go/) and Kyoto Encyclopedia of Genes and Genomes (KEGG) ^25^.

## 3. Results

### 3.1 Spatial transcriptomics performance on FFPE core needle biopsies of human kidney allografts

To understand whether FFPE core needle biopsies of human kidney allografts are feasible for spatial transcriptomics, we selected 8 cases based on histopathological and clinical features (Table 1), representing 4 diagnostic groups: 1) non-rejection conditions; 2) Active AMR; 3) Acute TCMR; 4) Chronic active AMR. The clinical diagnosis is interpreted by our renal pathologists based on the 2018 Banff Criteria ^26^. We performed 10x Genomic Visium spatial transcriptomics on H&E sections and obtained a transcriptomic map composed of “spots” (55 μm in diameter) with their own unique RNA expression signatures. The spots were localized on the kidney histological image with barcodes. We then used segmentation analysis to acquire 4-7 unsupervised clusters in each diagnostic category (Fig. 1). Visualization analysis based on the UMAP algorithm demonstrated the clusters with annotated labels, which were obtained based on histopathologic features and known marker genes associated with kidney structures (Fig. 1) ^27^. Due to the size of the spots employed in spatial transcriptomics, the cell densities associated with each barcode are variable, ranging from 1 to 10 per spot. As a result, some clusters showed admixed contribution of different cellular markers, such as tubular markers.

**Figure 1.**
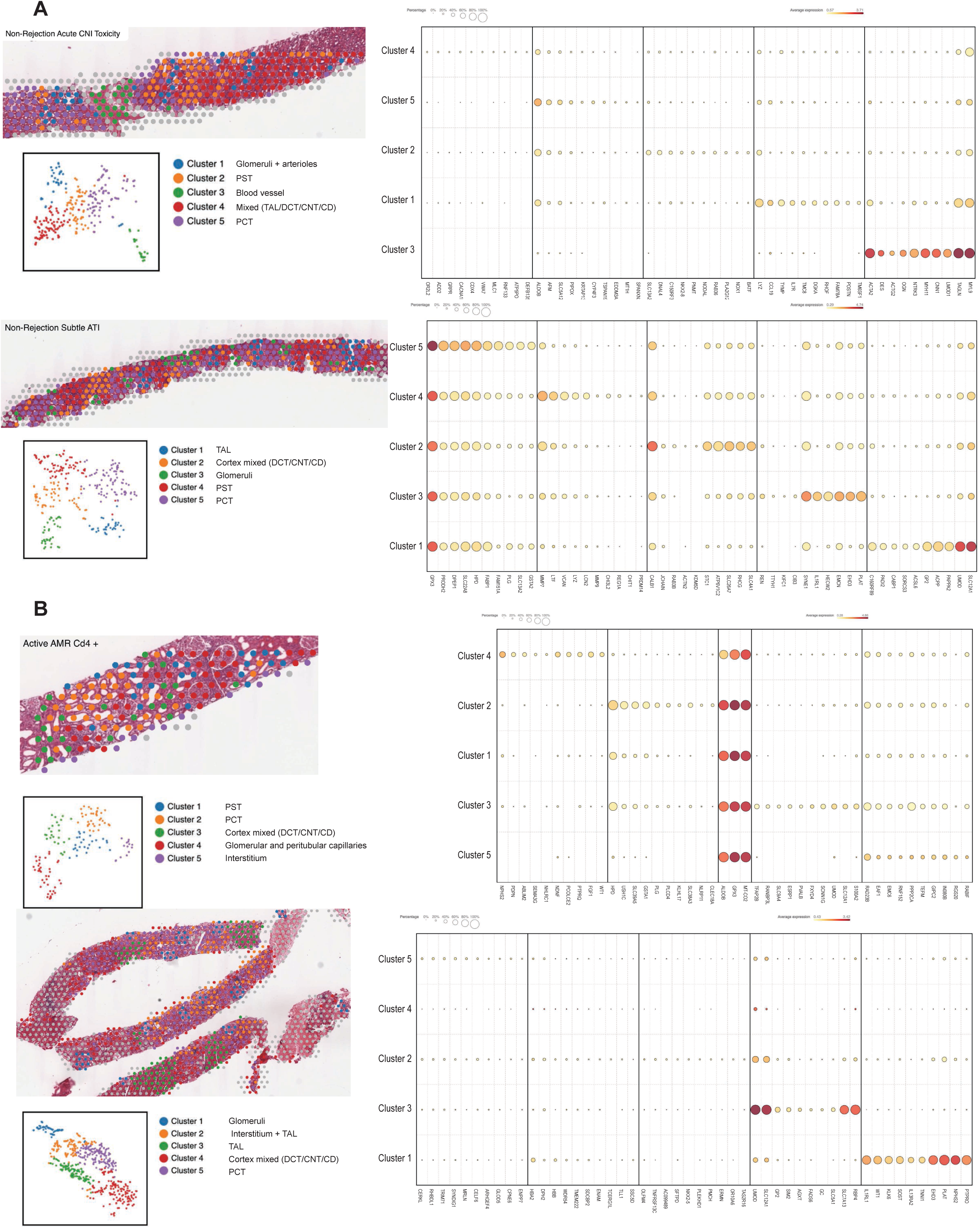
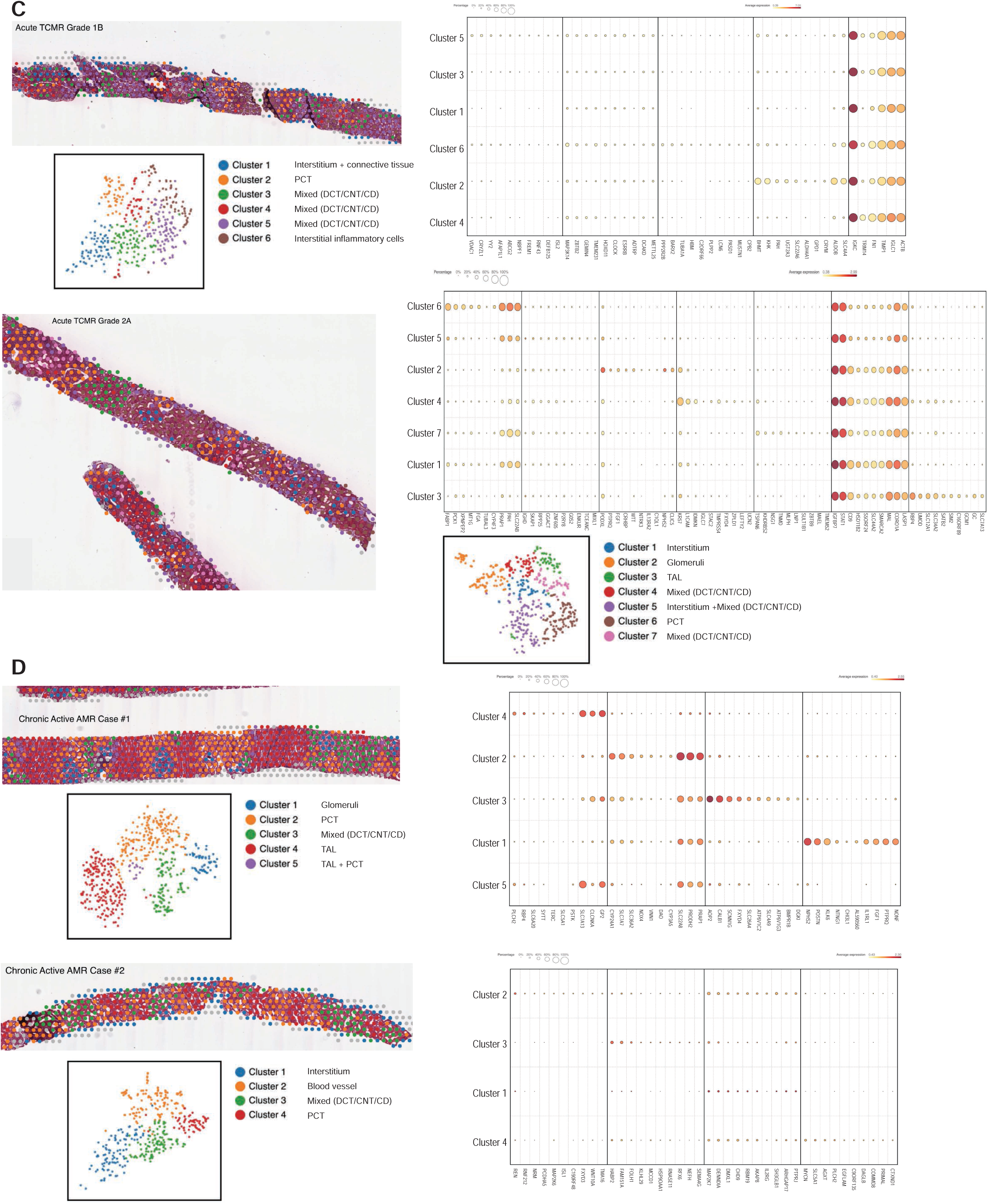
Spatial transcriptomics can be successfully performed on FFPE core needle biopsies of human kidney allografts. Transcriptomic map composed of “spots” (55 μm in diameter) with unique RNA expression signatures. The spots were localized on the kidney histologic image with barcodes. Four to seven unsupervised clusters in each case were shown by segmentation analysis and visualized by Uniform manifold approximation and projection (UMAP) algorithm. The cluster annotation was based on histopathologic features and known gene expression markers associated with kidney structures. CNI: acute calcineurin inhibitors; ATI: toxicity acute tubular injury; AMR: active antibody mediated rejection; TCMR: cell mediated rejection; PST: proximal straight tubule; PCT: proximal convoluted tubules; TAL: thick ascending limb; DCT: distal convoluted tubules; CNT: connecting tubules; CD: collecting duct.

**Table 1:** Histopathological and Clinical Features of Cases.

| Case # | 1 | 2 | 3 | 4 | 5 | 6 | 7 | 8 |
| --- | --- | --- | --- | --- | --- | --- | --- | --- |
| Diagnostic category | Non-rejection |  | Active AMR |  | Acute TCMR |  | Chronic active AMR |  |
| Pathologic diagnosis | Acute CNI toxicity | Subtle ATI | C4d-positive active AMR | C4d-negative active AMR | Acute TCMR, grade 1B, plasma cell rich | Acute TCMR, grade 2A | Chronic active AMR (Case #1) | Chronic active AMR (Case #2) |
| Banff Scores | t1, i0, v0, g0, ptc0, ci0, ct0, cg0, ti0, i-IFTA0, pvl0, C4d0 | t0, i0, v0, g0, ptc0, ci0, ct0, cg0, ti0, i-IFTA0, pvl0, C4d0 | t1, i1, v1, g2, ptc2, ci0, ct0, cg0, ti1, i-IFTA0, pvl0, C4d3 | t0, i0, v0, g2, ptc2, ci0, ct0, cg0, ti0, i-IFTA0, pvl0, C4d1 | t3, i3, v0, g0, ptc0, ci0, ct0, cg0, ti3, i-IFTA0, pvl0, C4d0 | t3, i2, v1, g0, ptc2, ci0, ct0, cg0, ti2, i-IFTA0, pvl0, C4d0 | t0, i0, v0, g2, ptc0, ci0, ct0, cg1b, ti0, i-IFTA0, pvl0, C4d2 | t0, i0, v0, g1, ptc1, ci0, ct0, cg2, ti0, i-IFTA0, pvl0, C4d2 |
| Age | 41 | 58 | 53 | 31 | 28 | 36 | 41 | 49 |
| Cause of ESKD | Hepato-renal syndrome | Diabetes | Unknown | Hypoplastic kidney | Unknown | Unknown | Unknown | Unknown |
| SCr (mg/dL) | 1.5 | 1.9 | 6.1 | 1.5 | 18 | 10 | 1.36 | 1.4 |
| DSA | Positive | Negative | Positive | Positive | Positive | Positive | Negative | Positive |
| Graft Function | DGF | DGF | Normal | Normal | Normal | Normal | Normal | Normal |
| Immuno-suppression Regimen | CSA, MMF, PRDL | TAC, MMF, PRDL | TAC, MMF, PRDL | TAC, MMF, PRDL | TAC, MMF, PRDL | TAC, MMF, PRDL | TAC, MMF, PRDL | TAC, MMF, PRDL |

CNI: acute calcineurin inhibitors; AMR: active antibody mediated rejection; ATI: toxicity acute tubular injury; Banff Score: tubulitis (t), interstitial inflammation in non-scarred areas (i), intimal arteritis (v), glomerulitis (g), peritubular capillaritis (ptc), interstitial fibrosis (ci), tubular atrophy (ct), glomerular basement membrane double contours (cg), total inflammation (ti), inflammation in the area of IFTA (i-IFTA), polyomavirus load (pvl); CSA: Cyclosporine A; DGF: delayed graft function; DSA, donor specific antibody; ESKD, end-stage kidney disease; IFTA, interstitial fibrosis and tubular atrophy; MMF: mycophenolate mofetil; PRDL: prednisolone; Scr: Serum creatine; TAC: Tacrolimus; TCMR: cell mediated rejection.

### 3.2 Different rejection types reveal distinct transcriptomic signatures

To identify DEGs in each of the rejection types with respect to non-rejection conditions, we used the Venice algorithm. Hierarchical clustering of the DEGs revealed distinct gene expression pattens among 8 cases (Fig. 2). Similar transcriptomic profiles patterns were observed among two cases in the same diagnostic groups (non-rejection cases, acute TCMR and chronic active AMR), except for the active AMR group. The C4d-positive active AMR case demonstrated significantly different transcriptomic signatures compared to the C4d-negative active AMR case, despite both being positive for donor-specific antibodies (DSA). Moreover, C4d-negative active AMR case showed a closer pattern to chronic active AMR cases. Furthermore, chronic active AMR cases shared some overlapping features with acute TCMR, which is consistent with recent study published by Shah et, al^28^. These results demonstrated that the transcriptomic signatures from FFPE core needle biopsy tissues have the potential to aid in distinguishing between different types of rejection and may also enable further subclassification of AMR.

**Figure 2.**
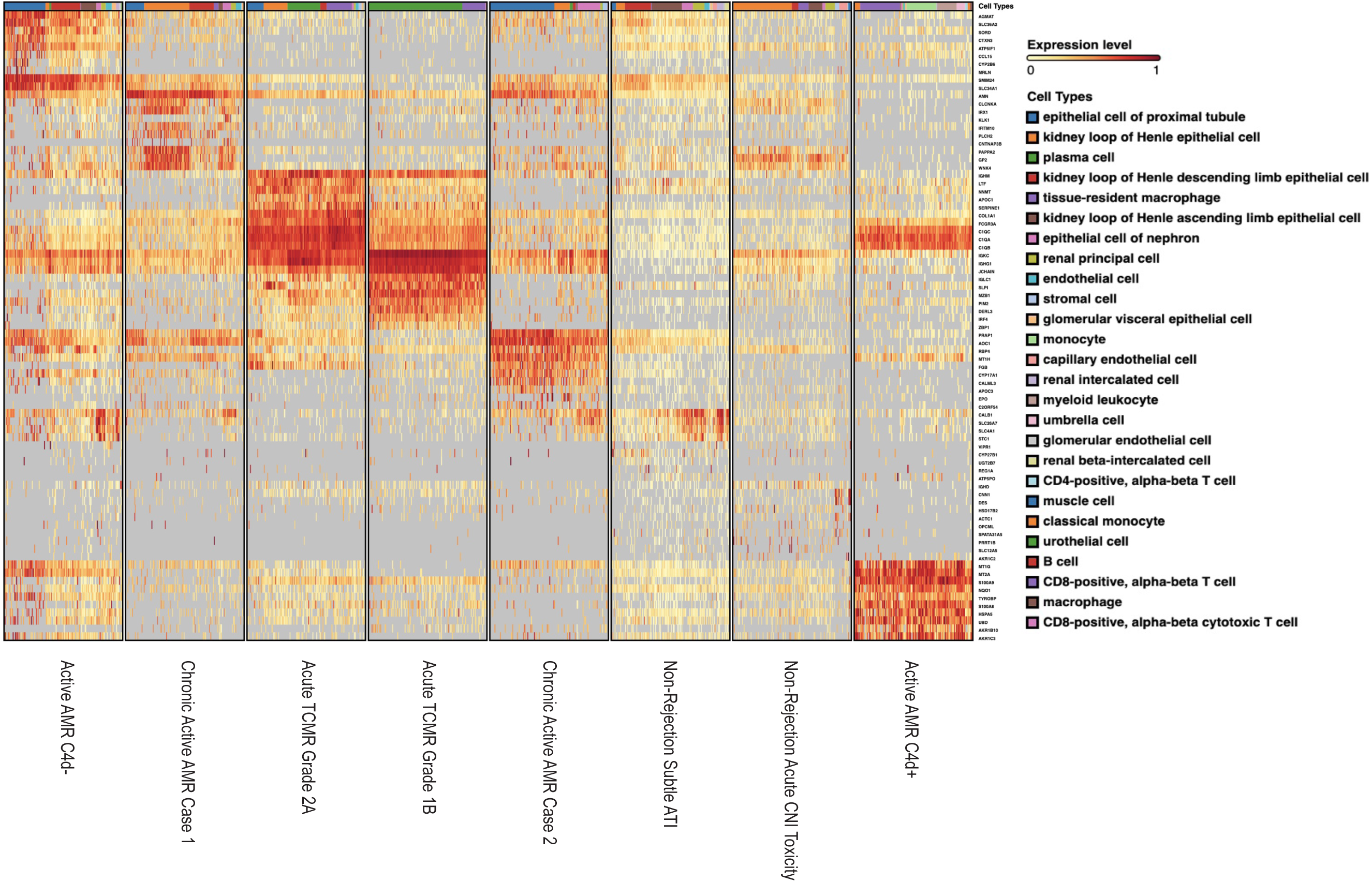
Different rejection types reveal distinct transcriptomic signatures. The heatmap, generated on the BioTuring platform, displayed differentially expressed genes (DEGs) organized via a dendrogram that illustrated the hierarchical relationships between cases. Color intensity represented gene expression levels, with red shades indicating higher expression and yellow shades indicating lower expression. The hierarchical clustering of rows (genes) and columns (cases) illustrated gene expression differences among four different diagnostic groups. These conditions exhibited distinct gene expression patterns, except for active AMR which demonstrated significantly different transcriptomic signatures between C4d-negative and C4d-positive active AMR cases. Chronic active AMR shared some overlapping features with acute TCMR. The dendrogram provided a visual representation of the genetic similarity and divergence among the studied cases.

To evaluate the concordance between DEG derived from our FFPE tissue transcriptomic signatures with top transcripts associated with rejection by MMDX ^1,29^, we compared two gene sets and observed some overlapping between our FFPE tissue transcriptomic signatures associated with active AMR and acute TCMR, and the MMDX transcripts linked to universal rejection (Halloran *et al*. 2017 paper) (Fig. 3A and 3B) ^1^. In addition, our FFPE tissue transcriptomic signatures associated with active AMR and acute TCMR showed some overlapping with the top 20 transcripts linked to AMR, TCMR, and injury- and rejection-associated transcripts as reported by Halloran *et al.* in their 2024 MMDX study (Fig. 3C and 3D) ^21^.

**Figure 3.**
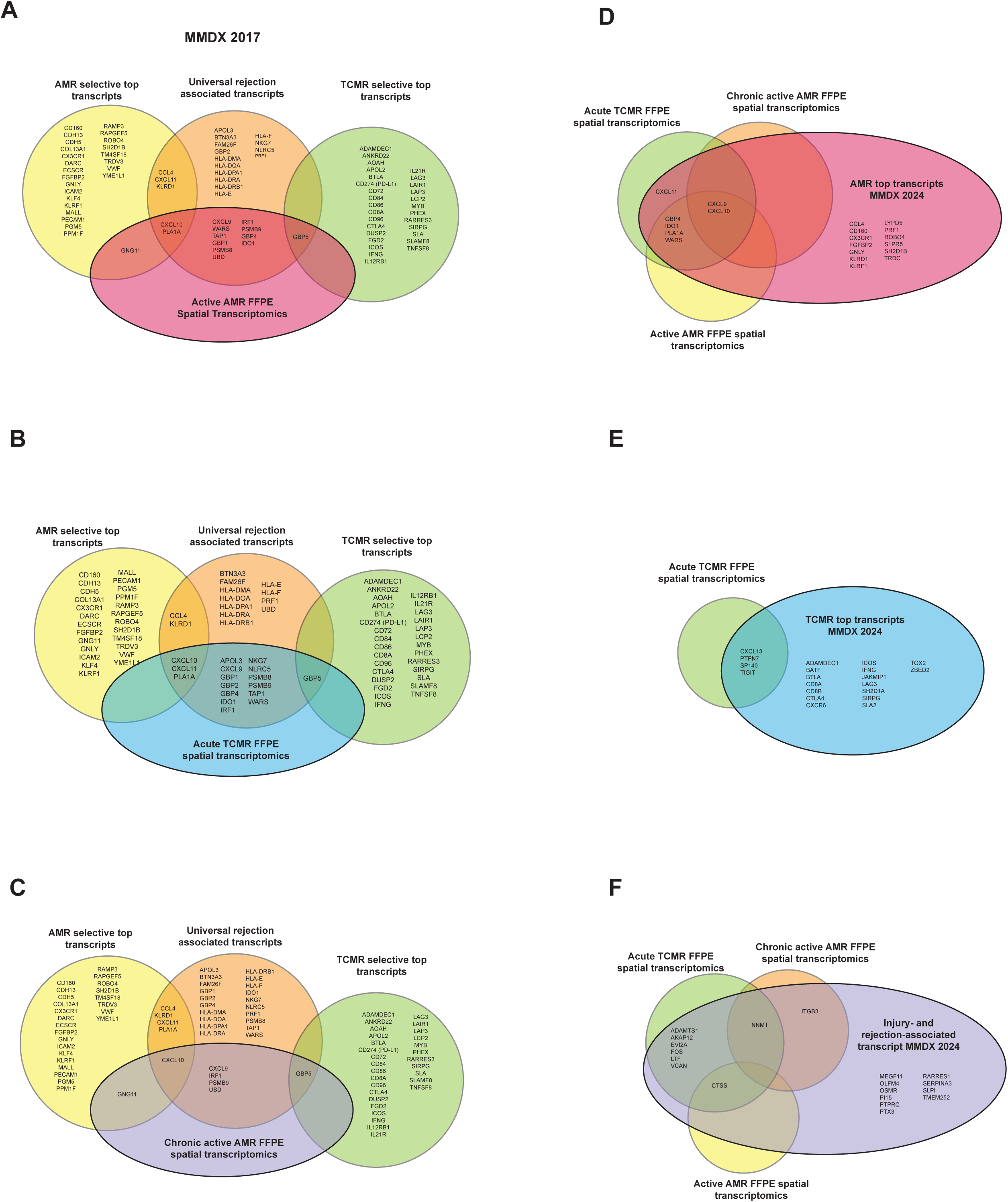
Overlap and distinctions between FFPE tissue transcriptomic signatures and published RNA signatures from bulk transcriptome microarrays. The Venn diagram illustrated the overlap and distinctions between our identified FFPE tissue transcriptomic signatures of active AMR (A), acute TCMR (B), chronic active AMR (C) and previously published top 30 transcripts associated with AMR-selected, TCMR-selected and universal rejection by MMDX (Halloran *et al*. 2017 paper)^1^. In addition, Venn diagram illustrated the overlap and distinctions between our identified FFPE tissue transcriptomic signatures of active AMR (D), acute TCMR (E), and chronic active AMR (F) and previously published top 20 transcripts associated AMR, TCMR, and injury- and rejection-associated transcript by MMDX (Halloran *et al*. 2024 paper), respectively ^29^.

Furthermore, our analysis of the top 30 transcriptomic signatures in FFPE tissue from rejection groups (Table 2) revealed additional important genes that are associated with transplant rejection. For example, in C4d-positive active AMR case, *S100A8* and *S100A9* were significantly upregulated. These calcium-binding proteins, primarily expressed in monocytes, play a crucial role in kidney transplant rejections, and high expression levels of S100A8 and S100A9 in myeloid cells during kidney transplant rejections have been linked to favorable outcomes ^30^. In acute TCMR, the expression of *FCGR3A* gene, which encodes the Fc gamma receptor IIIA (FcγR IIIA or CD16), was significantly increased, with its specific role to be elaborated upon later. Additionally, Interferon Regulatory Factor 4 (*IRF4*) was significantly upregulated. Similar to *IRF1*, this transcription factor is critical for immune regulation, particularly in T and B cells, and plays a significant role in transplant rejection by regulating genes involved in inflammation and lymphocyte activation ^31^. Moreover, the expression of complement component C3 was significantly increased. C3, part of the complement system that is frequently activated in acute AMR ^31^, was also significantly increased in acute TCMR.

**Table 2:** Top 30 genes (sorted by Log2 fold change (Log2FC) value) selectively increased in biopsies with active AMR, acute TCMR and chronic active AMR.

| Active AMR, C4d- |  |  | Active AMR, C4d+ |  |  | Acute TCMR, grade 1B |  |  | Acute TCMR, grade 2A |  |  | Chronic active AMR, case #1 |  |  | Chronic active AMR, case #2 |  |  |
| --- | --- | --- | --- | --- | --- | --- | --- | --- | --- | --- | --- | --- | --- | --- | --- | --- | --- |
| Genes | p-value | Log2 FC | Genes | p-value | Log2 FC | Genes | p-value | Log2 FC | Genes | p-value | Log2 FC | Genes | p-value | Log2 FC | Genes | p-value | Log2 FC |
| AGMAT | 5.0E-131 | 2.91 | AKR1B10 | 3.4E-64 | 5.34 | IGLC1 | 2.4E-228 | 3.96 | APOC1 | 3.5E-147 | 3.81 | PAPPA2 | 1.1E-78 | 2.69 | CALML3 | 4.6E-44 | 3.60 |
| CTXN3 | 1.4E-73 | 2.78 | **S100A8** | 2.9E-88 | 4.67 | DERL3 | 2.6E-131 | 3.9 | LTF | 3.8E-205 | 3.69 | IFITM10 | 3.5E-43 | 2.54 | EPO | 1.3E-03 | 2.80 |
| SMIM24 | 2.7E-202 | 2.7 | UBD | 2.2E-82 | 4.57 | IGKC | 3.9E-252 | 3.65 | COL1A1 | 1.9E-265 | 3.63 | PLCH2 | 6.2E-29 | 2.42 | APOC3 | 6.1E-33 | 2.62 |
| CYP2B6 | 3.7E-22 | 2.69 | AKR1C3 | 1.9E-69 | 4.46 | MZB1 | 2.5E-192 | 3.56 | NNMT | 1.9E-200 | 3.28 | AMN | 1.7E-178 | 2.40 | CYP17A1 | 2.9E-52 | 2.56 |
| SORD | 8.9E-68 | 2.63 | **S100A9** | 2.8E-93 | 4.34 | SLPI | 1.1E-166 | 3.41 | SERPINE1 | 1.8E-146 | 3.08 | IRX1 | 3.1E-52 | 2.35 | PRAP1 | 1.8E-101 | 2.51 |
| SLC34A1 | 2.4E-115 | 2.45 | HSPA5 | 3.8E-69 | 4.25 | JCHAIN | 3.7E-241 | 3.24 | C1QC | 3.1E-267 | 3.07 | CNTNAP3B | 4.3E-04 | 2.32 | MT1H | 1.5E-89 | 2.46 |
| ATP5IF1 | 8.8E-76 | 2.44 | MT2A | 2.6E-90 | 4.09 | **IRF4** | 2.2E-101 | 3.05 | **FCGR3A** | 6.5E-224 | 2.97 | KLK1 | 2.3E-13 | 2.23 | RBP4 | 9.9E-56 | 2.29 |
| SLC36A2 | 5.9E-74 | 2.41 | NQO1 | 5.7E-78 | 3.98 | ZBP1 | 6.7E-53 | 3.03 | IGHM | 4.0E-199 | 2.94 | GP2 | 7.5E-53 | 2.18 | FGB | 1.3E-58 | 2.27 |
| MRLN | 4.0E-07 | 2.4 | MT1G | 5.3E-59 | 3.94 | IGHG1 | 2.0E-215 | 2.74 | C1QA | 1.9E-257 | 2.92 | CLCNKA | 1.2E-49 | 2.13 | C2ORF54 | 4.8E-09 | 2.24 |
| CCL15 | 2.1E-20 | 2.39 | TYROBP | 2.1E-76 | 3.67 | PIM2 | 1.6E-123 | 2.71 | C1QB | 1.3E-253 | 2.87 | WNK4 | 1.0E-65 | 2.10 | AOC1 | 7.6E-110 | 2.23 |
| FAM151A | 7.7E-55 | 2.38 | CYP4F11 | 2.9E-45 | 3.57 | ANGPTL4 | 1.5E-37 | 2.68 | CD163 | 5.2E-219 | 2.78 | MROH7 | 1.5E-29 | 1.99 | SERPINA5 | 7.4E-66 | 2.18 |
| MT1X | 3.7E-111 | 2.35 | PDIA4 | 6.2E-51 | 3.48 | XBP1 | 8.0E-178 | 2.64 | FN1 | 9.9E-218 | 2.73 | SLC9A3 | 1.3E-85 | 1.90 | SCNN1G | 5.6E-34 | 1.97 |
| MDH1 | 5.3E-54 | 2.35 | CCL13 | 6.3E-29 | 3.48 | TXNDC5 | 1.4E-213 | 2.62 | **C3** | 3.4E-165 | 2.62 | C16ORF89 | 8.4E-34 | 1.87 | VTN | 3.5E-05 | 1.93 |
| HBA2 | 1.8E-04 | 2.31 | GNG5 | 8.6E-13 | 3.4 | CD79A | 3.7E-61 | 2.59 | COL1A2 | 7.8E-221 | 2.61 | CLCNKB | 2.7E-37 | 1.85 | NCAM2 | 1.6E-14 | 1.86 |
| NDUFS8 | 1.2E-28 | 2.28 | GCLM | 4.3E-58 | 3.3 | BLK | 7.3E-27 | 2.56 | CLDN1 | 5.5E-99 | 2.56 | FMO4 | 7.3E-95 | 1.84 | TRIM50 | 2.2E-06 | 1.85 |
| PCK1 | 2.7E-101 | 2.28 | PRDX1 | 4.6E-70 | 3.26 | FCRL5 | 1.3E-56 | 2.56 | VSIG4 | 3.4E-164 | 2.56 | ACPP | 1.2E-46 | 1.82 | C2ORF40 | 1.9E-27 | 1.85 |
| GATM | 7.0E-183 | 2.24 | HIST1H2BB | 1.1E-10 | 3.25 | PDK1 | 5.7E-112 | 2.55 | IFITM3 | 2.4E-247 | 2.42 | AGXT | 4.0E-08 | 1.82 | GSTA1 | 7.6E-93 | 1.83 |
| C1ORF35 | 9.8E-28 | 2.23 | FCER1G | 1.1E-68 | 3.19 | SPAG4 | 2.6E-32 | 2.53 | SLC1A3 | 1.0E-67 | 2.38 | HSPA1A | 8.7E-97 | 1.78 | RHCG | 5.2E-33 | 1.83 |
| MIOX | 9.8E-135 | 2.22 | ATP5PO | 2.9E-07 | 3.08 | TNFRSF17 | 5.5E-44 | 2.52 | IFITM1 | 4.4E-105 | 2.33 | AQP7 | 2.5E-57 | 1.76 | PCDH10 | 1.1E-21 | 1.82 |
| XYLB | 9.1E- | 2.21 | NOP10 | 1.9E- | 3 | LAX1 | 9.3E- | 2.5 | MS4A4A | 2.7E- | 2.32 | ETV4 | 4.2E- | 1.73 | TLDC2 | 4.3E- | 1.80 |
|  | 30 |  |  | 28 |  |  | 46 |  |  | 109 |  |  | 21 |  |  | 06 |  |
| PTMS | 1.4E-44 | 2.19 | CYBB | 2.8E-75 | 2.99 | ZNF215 | 2.5E-31 | 2.42 | ADAMTS2 | 8.4E-54 | 2.31 | ACSL6 | 5.0E-12 | 1.71 | LEAP2 | 6.0E-08 | 1.80 |
| SLC34A3 | 1.6E-29 | 2.18 | GMFG | 1.4E-25 | 2.98 | MS4A1 | 2.2E-39 | 2.38 | FCGR1B | 3.1E-78 | 2.31 | EFHD1 | 1.0E-74 | 1.68 | BSG | 1.2E-131 | 1.79 |
| TMBIM6 | 9.7E-16 | 2.17 | CALR | 9.7E-71 | 2.95 | PRDM1 | 1.7E-79 | 2.36 | CDH6 | 4.9E-110 | 2.30 | GPT | 3.0E-51 | 1.67 | TRIM6 | 8.9E-44 | 1.78 |
| FMO1 | 3.5E-68 | 2.12 | HINT1 | 9.2E-41 | 2.93 | TENT5C | 9.5E-108 | 2.34 | MVP | 6.3E-138 | 2.28 | EPHB3 | 1.4E-25 | 1.67 | FABP1 | 3.9E-58 | 1.78 |
| UQCRB | 1.2E-69 | 2.09 | MARCO | 1.0E-28 | 2.89 | POU2AF1 | 4.2E-46 | 2.31 | LCN2 | 3.1E-24 | 2.23 | CAPN12 | 1.8E-27 | 1.65 | CTSV | 1.3E-12 | 1.76 |
| EMCN | 5.4E-20 | 2.09 | VNN1 | 1.8E-38 | 2.88 | IGHA1 | 2.9E-161 | 2.28 | CCL4L2 | 1.1E-39 | 2.21 | AL845331 | 2.8E-19 | 1.64 | GRP | 2.7E-02 | 1.75 |
| ZNF787 | 4.4E-06 | 2.07 | EIF1 | 5.9E-67 | 2.87 | FPR1 | 9.7E-17 | 2.27 | CXCL11 | 6.8E-110 | 2.19 | GHDC | 1.3E-56 | 1.62 | GSTA2 | 1.6E-23 | 1.73 |
| C3ORF85 | 2.3E-06 | 2.06 | NDUFB4 | 1.2E-29 | 2.87 | ANKDD1A | 3.2E-14 | 2.17 | CXCL13 | 2.7E-45 | 2.18 | GGACT | 7.4E-53 | 1.61 | OLR1 | 4.1E-16 | 1.67 |
| HBB | 5.7E-10 | 2.05 | PSMA4 | 3.0E-55 | 2.87 | ZNF534 | 1.9E-09 | 2.14 | FKBP5 | 4.9E-132 | 2.17 | MFSD3 | 2.5E-50 | 1.61 | PRELP | 1.7E-17 | 1.67 |

The genes highlighted in bold are those that play a role in acute rejection but were not listed among the top transcripts in the MMDX study.

### 3.3 Distinct subclusters of monocytes/macrophages exhibiting high FCGR3A expression are identified in acute rejection groups

Acute rejection poses a significant threat to allograft survival. It is crucial to identify the specific cell populations that play key roles in various forms of acute rejection. Understanding these cellular dynamics is essential for developing potential innovative targeted therapies and improving long-term transplant outcomes. Therefore, we performed a joint visualization of spots in all cases using t-SNE dimension reduction method. The cell type composition of each case (Fig. 4A) was generated by referencing the expression profiles of 10X Visium bins against a published meta-database of characterized kidney cells using BioTuring. Acute TCMR cases demonstrated a prominent tissue-resident macrophage population (Fig. 4B). These tissue-resident macrophages (markers: CD68 and CD163) exhibited high expression of *FCGR3A* (Fig. 4C). The “Monocyte category” includes classical (FCGR3A- and CD14+), intermediate monocytes (FCGR3A+ and CD14+) and non-classical monocytes (FCGR3A+ and CD14-), while the “classical monocyte” category specifically represents the classical monocytes^32^. The analysis revealed that the C4d-positive active AMR case showed a significant population of non-classical and intermediate monocytes, which was the highest among and significantly different from all other cases (Fig. 4B). This distinct subcluster of monocytes (markers: CD14 and CD68) demonstrated a high *FCGR3A* expression (Fig. 4D). *FCGR3A* is involved in cellular cytotoxicity and is thought to play a significant role in acute rejection ^6,33^. Spatial transcriptomics data analysis of *FCGR3A* expression using UMAP visualization for each case is shown in Supplementary Figure 1.

**Figure 4:**
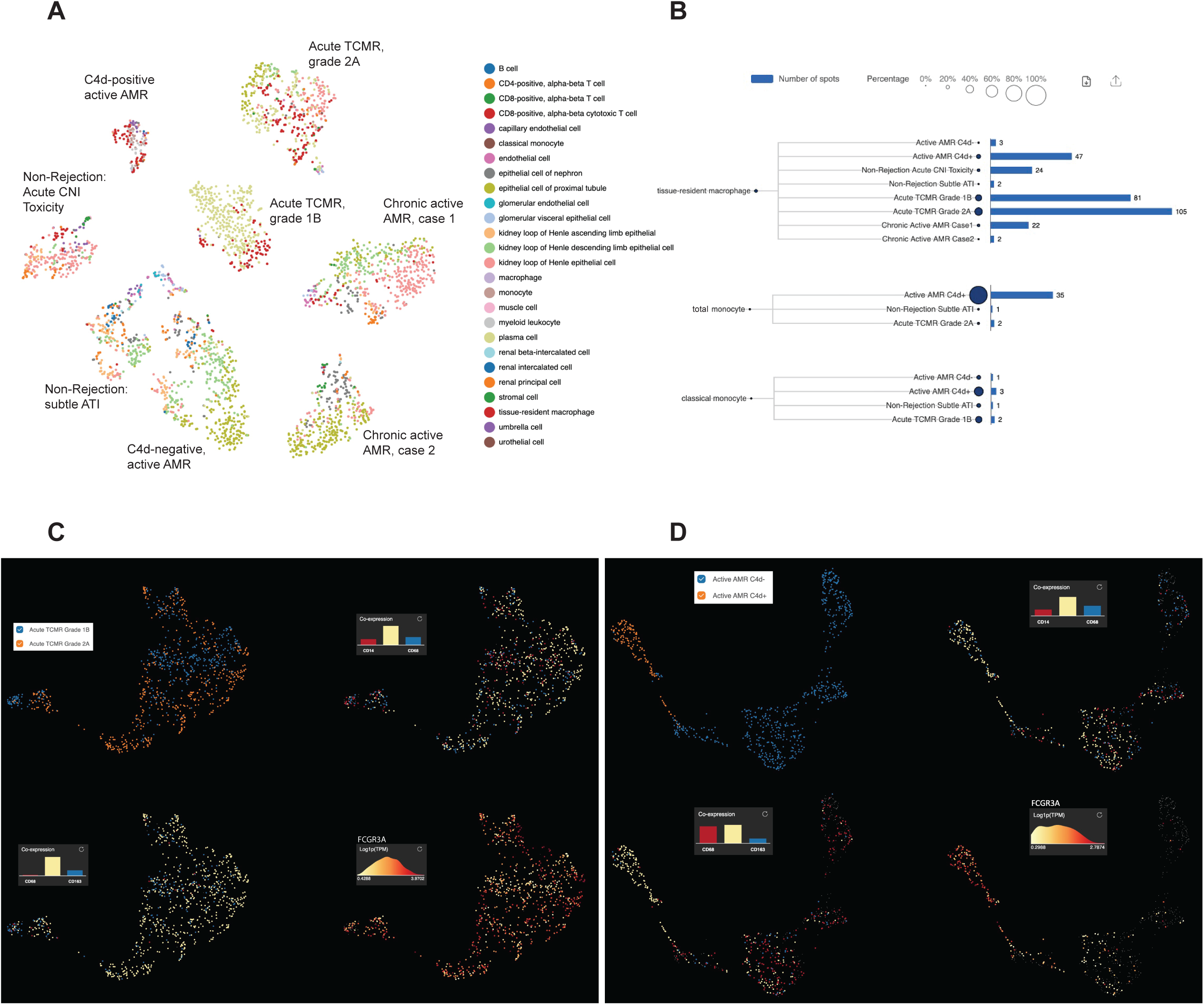
Distinct subclusters of monocytes/macrophages are identified in acute rejection groups with high *FCGR3A* expression. The t-distributed stochastic neighbor embedding (t-SNE) dimension reduction and cell composition of each case was shown in (A). The number of spots and percentage of macrophages, total monocytes and classical monocytes were illustrated in (B). Prominent tissue-resident macrophage populations were identified in acute TCMR cases, and a significant population of non-classical and intermediate monocytes (total monocytes minus classic monocytes) was identified in C4d-positive active AMR case. UMAP analysis of acute TCMR grade 1B (blue) and grade 2A (orange) was shown in (C). The clusters are overlaid with expression markers for monocytes (CD14 and CD68), macrophage (CD68 and CD163) and Fc gamma receptor IIIA (*FCGR3A*). It revealed distinct macrophage/monocytes subclusters exhibiting high expression of *FCGR3A* were evident. Similarly, UMAP analysis comparing C4d-negative (blue) and C4d-positive (orange) was shown in (D). These clusters were also overlaid by monocytes and macrophage markers, as well as *FCGR3A,* which revealed distinct macrophage/monocytes subclusters with high expression of *FCGR3A*.

### 3.4 Spatial distribution of monocyte/macrophage subclusters with high FCGR3A expression corresponded to the characteristic histopathological features in acute rejection groups

To identify the spatial locations of these distinct monocyte/macrophage subclusters, the expression of monocyte/macrophage markers and *FCGR3A* was mapped onto the biopsy H&E images using Loupe Browser (Fig. 5). In C4d-positive AMR, clusters over representative areas of peritubular capillaritis (PTCitis) and glomerulitis showed enrichment in both monocyte/macrophage markers and *FCGR3A* expression (Fig. 5A-B). In acute TCMR, both grade 1B (Fig. 5C-D) and grade 2A (Fig. 5E-F) cases demonstrated enrichment of monocyte/macrophage markers and *FCGR3A* expression in clusters over representative areas of tubulitis and interstitial inflammation. Additionally, inflammatory cells in the intimal arteritis (V1 lesion) of the acute TCMR grade 2A case exhibited high co-expression of monocyte/macrophage markers and *FCGR3A* (Fig. 5E-F).

**Figure 5:**
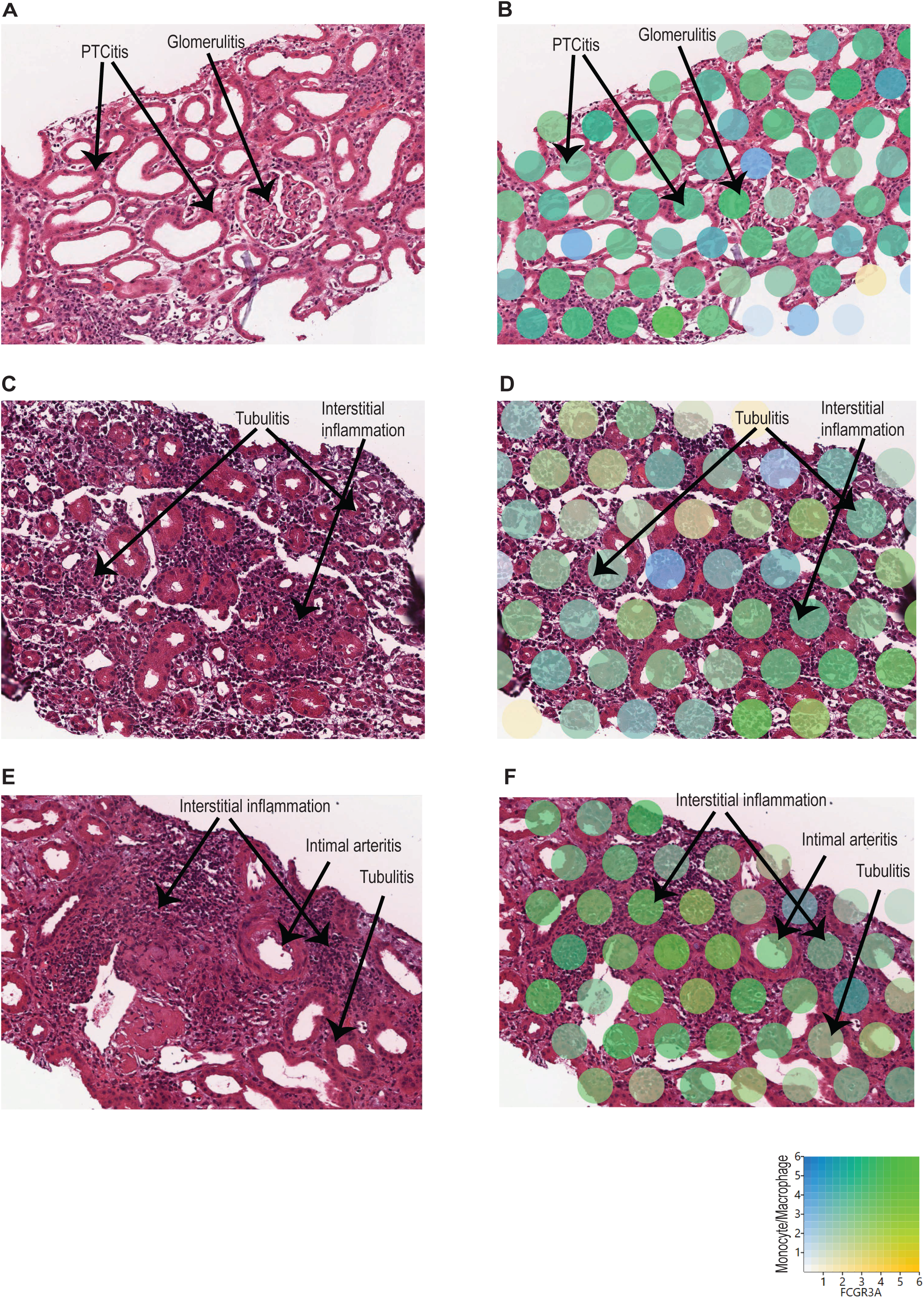
Spatial location of monocyte/macrophage subclusters with high *FCGR3A* expression. The expression of monocyte/macrophage markers (blue) and *FCGR3A* (yellow) was mapped onto the biopsy H&E images using Loupe Browser, using Log2 as scale value. Co-expression is indicated in green. A-B) C4d-positive AMR: Clusters over representative areas of peritubular capillaritis (PTCitis) and glomerulitis showed enrichment in both monocyte/macrophage markers and *FCGR3A* expression. C-D) Acute TCMR, grade 1B: Enrichment of monocyte/macrophage markers and *FCGR3A* expression in clusters over representative areas of tubulitis and interstitial inflammation. E-F) Acute TCMR, grade 2A: Enrichment of monocyte/macrophage markers and *FCGR3A* expression in clusters over representative areas of tubulitis and interstitial inflammation. In addition, high co-expression of monocyte/macrophage markers and *FCGR3A* in inflammatory cells within the intimal arteritis (V1 lesion).

### 3.5 Functional pathway and gene network analysis

To identify enriched functional pathway associated with DEG, we performed functional pathway analysis of the DEGs using GO enrichment analysis and KEGG analysis ^25^. GO analysis revealed top perturbed GO biological process pathways enriched in all rejection groups, with key pathways associated with metabolic changes in trained immunity (Fig. 6A-D). For instance, carboxylic acid catabolic, amino acid and fatty acid metabolic process pathways were upregulated in C4d-negative active AMR (Fig. 6A). Intermediates from these process can enter glycolysis and the tricarboxylic acid (TCA) cycle, linking these pathways together ^34^. In chronic active AMR, there was an increase in aerobic glycolysis and mitochondrial oxidative metabolism (such as oxidative phosphorylation, respiratory electron transport chain, and Adenosine triphosphate (ATP) synthesis) (Fig. 6D). In contrast to C4d-negative active AMR and chronic active AMR, we observed several key immune-related pathways in C4d-positive active AMR (Fig. 6B). These included pathways involved in activating and regulating immune responses, as well as those regulating innate immune responses and NF-kappa B signaling. These findings parallelled our observations in acute TCMR, where we also identified upregulation of pathways associated with mononuclear cells (lymphocytes and monocytes/macrophages) differentiation, immune response-activating signaling pathways, phagocytosis, and regulation of innate immune response (Fig. 6C).

**Figure 6:**
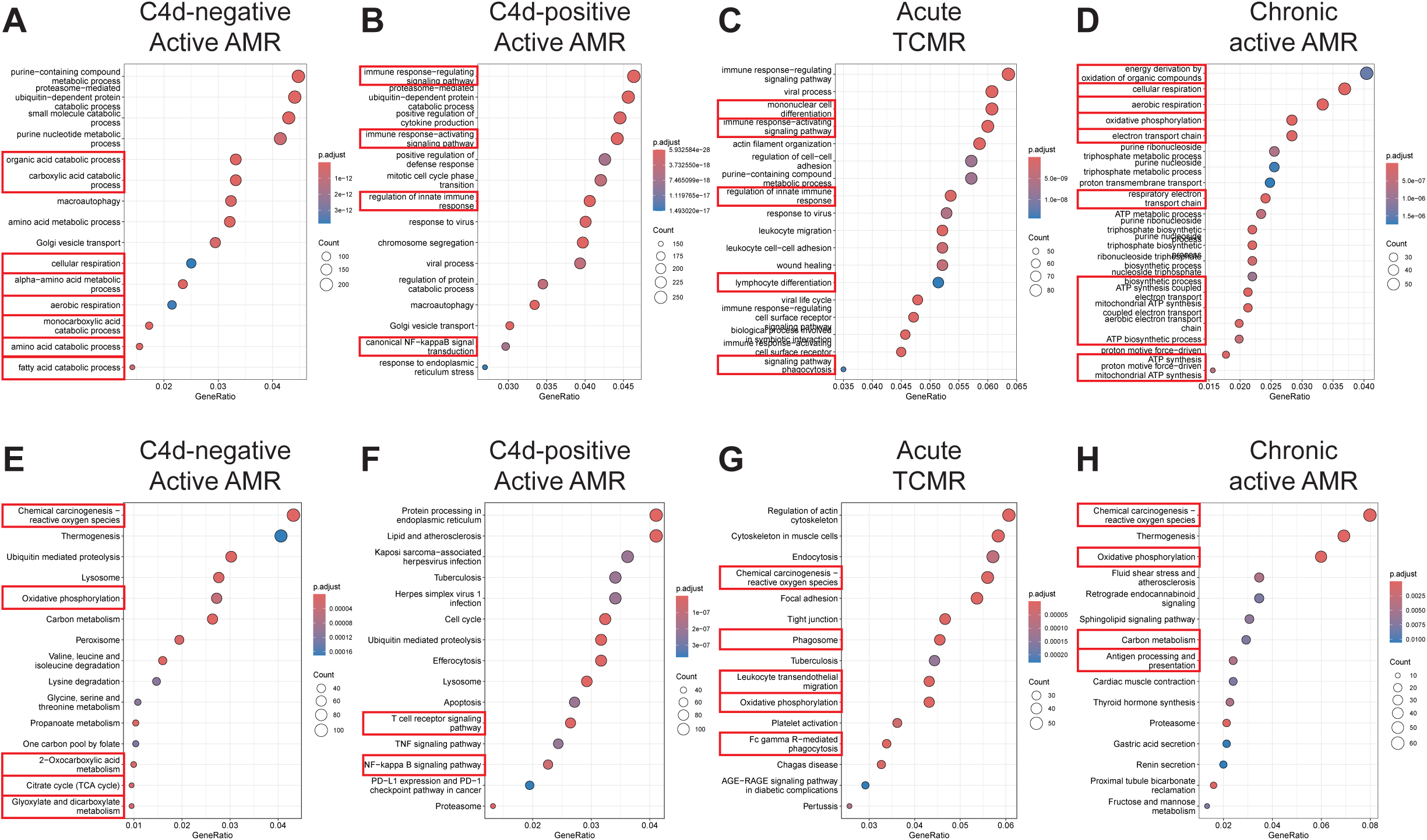
Functional pathway and gene network analysis. A-D) Gene Ontology (GO) Enrichment Analysis. Key pathways (highlighted with red rectangles) associated with metabolic changes in trained immunity were upregulated in C4d-negative active AMR (A) and chronic active AMR (D). In C4d-positive active AMR (B) and acute TCMR (C), we observed upregulation of pathways related to activation and regulation of immune response including innate immunity. E-H) Kyoto Encyclopedia of Genes and Genomes (KEGG) Analysis. Key metabolic pathways (highlighted with red rectangles) aligned with the GO analysis in both C4d-negative active AMR (E) and chronic active AMR (H). In addition to GO analysis, KEGG analysis revealed upregulation of additional rejection-associated damage and macrophage response to transplant allografts pathways in TCMR (G). It also highlighted antigen processing and presentation pathways in chronic active AMR (H).

KEGG analysis supported the GO analysis findings, revealing similar upregulation of metabolic pathways in both C4d-negative active AMR and chronic active AMR (Fig 6E and 6H). Moreover, both conditions exhibited increased ROS production. In addition to the immune-related pathways identified in the GO analysis, KEGG analysis uncovered upregulation of additional rejection-associated damage and macrophage response to transplant allografts pathways in TCMR, including ROS production, leukocyte trans-endothelial migration and FcγR-mediated phagocytosis (Fig. 6G). Furthermore, KEGG analysis revealed upregulation of antigen processing and presentation pathways in chronic active AMR (Fig. 6H).

### 3.6 Upregulation of CD47 and SIPR***α*** in acute rejection

Innate allorecognition, which allows innate immune cells to discriminate between self and non-self, is one of the most important mechanisms of innate immune activation during acute transplant rejection ^35^. CD47, leukocyte immunoglobulin-like receptor A (LILRA), and signal-regulatory protein-α (SIPRα) are key markers associated with monocytes/macrophage activation and function in both transplant rejection and trained immunity within the innate alloantigen recognition pathway ^36^. SIPRα and MHC class I antigens are expressed on allografts and are recognized by CD47 and LILRA that are expressed on host monocytes, respectively. The UMAP visualization and violin plots of log2 fold changes illustrated significantly higher expression of *CD47* (*p*<0.05) and notably higher expression of *SIRP*α in C4d-positive active AMR. *CD47* and *SIPR*α expression are also upregulated in acute TCMR cases. However, this upregulation is not as pronounced as in the C4d-positive active AMR case (Fig. 7A and B). The interaction between *FCGR3A* and *LILRA* is believed to play important roles in monocytes/macrophage activation and function during transplant rejection ^36–38^, and we observed significant upregulation of *FCGR3A* in C4d-positive active AMR and acute TCMR cases (Fig. 7A and B). However, we did not observe *LILRA* expression upregulation among these cases.

**Figure 7:**
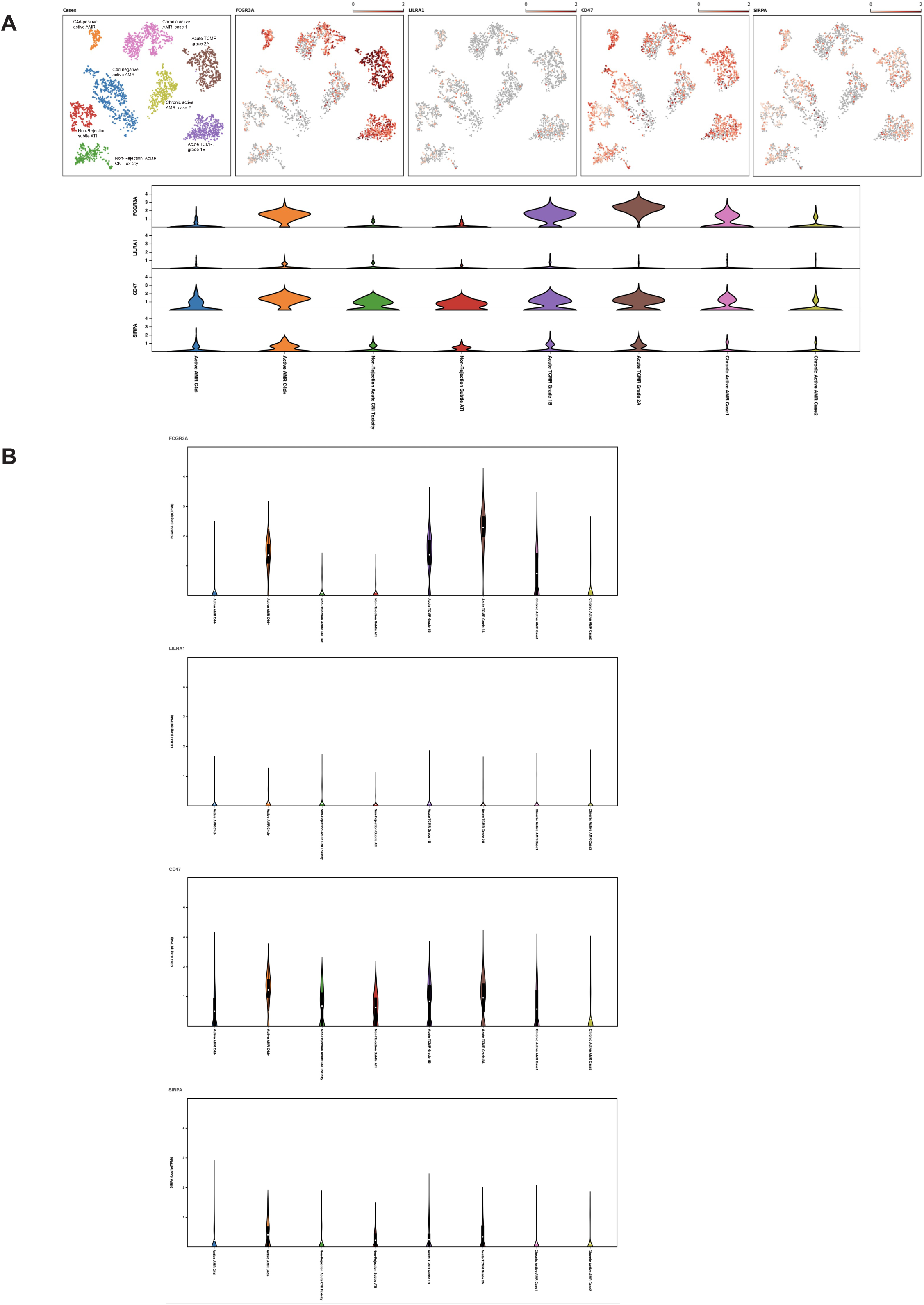
Upregulation of *CD47 and SIPR*α in acute rejection. A) The UMAP visualization, and violin plots of log2 fold changes (A and B) illustrated that *CD47* expression was significant higher and *SIRP*α expression was notably higher in C4d-positive active AMR case. *CD47* and signal-regulatory protein-α (*SIPR*α) expression were also upregulated in acute TCMR cases, but not as pronounced as in the C4d-positive active AMR case. *FCGR3A* was significantly upregulated in both C4d-positive active AMR and acute TCMR cases. However, we did not observe leukocyte immunoglobulin-like receptor A (*LILRA*) expression upregulation among these cases. The box within each violin plot (B) represents the interquartile range of the data, with the mean represented as a white dot.

### 3.7 Metabolic genes expression related to trained immunity

The expression of key metabolic gene markers across different groups for trained immunity, including the key genes involved in glycolysis and mitochondrial oxidative metabolism were depicted as bubble plot (Fig. 8). The bubble plot also included genes that encode metabolic intermediates, which are believed to induce epigenetic changes, such as fumarase (*FH*) gene and succinate dehydrogenase complex (*SDHA/SDHB/SDHC/SDHD*). This analysis revealed distinct expression patterns between groups experiencing rejection and those without rejection. Non-rejection conditions, such as acute CNI toxicity and subtle ATI, showed elevated activity in the mTOR pathway, glycolysis, and mitochondrial oxidative metabolism. In contrast, all rejection groups exhibited more pronounced elevations in glycolysis and mitochondrial oxidative metabolism activities than mTOR pathway activity. Notably, within glycolysis-related genes, Enolase 1 (*ENO*1) showed a significant increase in non-rejection conditions and C4d-negative active AMR, while Pyruvate kinase (*PKM*) was significantly elevated in acute TCMR groups and chronic active AMR. C4d-positive active AMR displayed significant increases in both genes. Additionally, clusters associated with acute TCMR and chronic active AMR showed evidence of increased levels of metabolic intermediates, *SDHA/SDHB*, which are thought to induce epigenetic changes.

**Figure 8:**
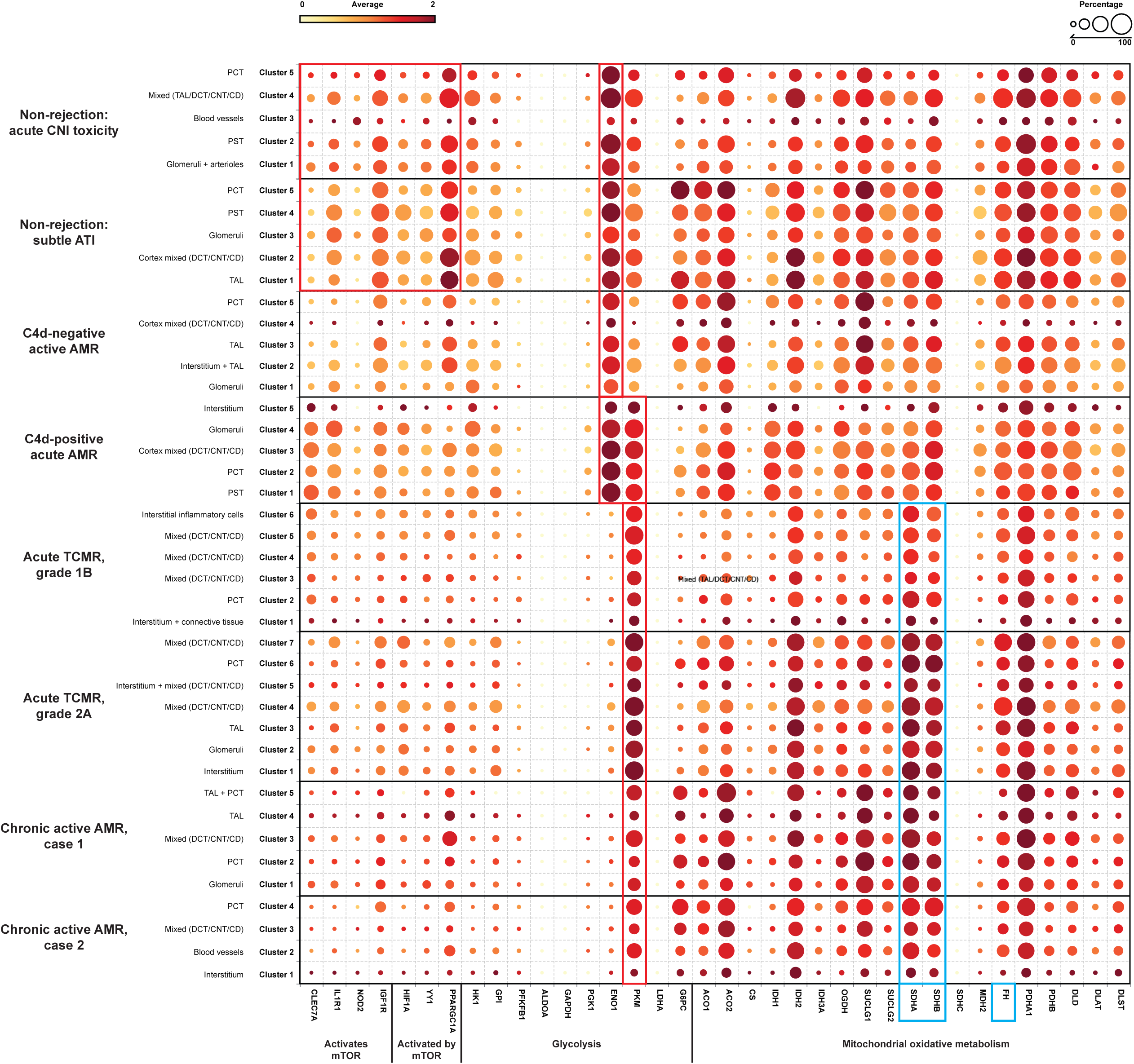
Metabolic genes expression related to trained immunity. A dotplot analysis of metabolic genes related to trained immunity revealed distinct patterns across non-rejection and rejection conditions. Non-rejection conditions (acute CNI toxicity and subtle ATI) showed increased activity in mTOR, glycolysis, and mitochondrial oxidative metabolism, while all rejection groups exhibited more pronounced glycolytic and oxidative metabolism. Notably, enolase 1 (*ENO*1) was elevated in non-rejection conditions and C4d-negative active AMR, while pyruvate kinase (*PKM*) was significantly increased in acute TCMR and chronic active AMR (red rectangle). C4d-positive active AMR showed significant increases in both genes (red rectangle). Acute TCMR and chronic active AMR clusters also displayed elevated levels of succinate dehydrogenase A/B (*SDHA/SDHB*), metabolic intermediates associated with epigenetic changes (blue rectangle). The dotplot includes genes: 1) genes activate mTOR pathway: *CLEC7A* (C-type lectin domain family 7 member A), *IL1R1* (Interleukin 1 Receptor Type 1), *NOD2* (Nucleotide Binding Oligomerization Domain Containing 2), *IGF1R* (Insulin Like Growth Factor 1 Receptor); 2) genes activated by mTOR pathway: *HIF1A* (Hypoxia-Inducible Factor 1-alpha), *YY1* (Yin Yang 1), *PPARGC1A* (Peroxisome proliferator-activated receptor-γ coactivator 1-α); 3) glycolysis: *HK*1 (Hexokinase 1), *GPI* (Glucose-6-phosphate isomerase), *PFKFB1* (6-Phosphofructo-2-Kinase/Fructose-2,6-Biphosphatase 1), *ALDOA* (Aldolase A), *GAPDH* (Glyceraldehyde-3-phosphate dehydrogenase), *PGK1* (Phosphoglycerate kinase 1), *ENO*1, *PKM*, *LDHA* (Lactate dehydrogenase A), *G6PC* (Glucose-6-phosphatase); 4) mitochondrial oxidative metabolism: *ACO1/ACO2* (Aconitase), *CS* (Citrate synthase), *IDH1/IDH2* (Isocitrate dehydrogenase), *OGDH* (α-ketoglutarate dehydrogenase), *SUCLG1/SUCLG2* (Succinyl-CoA ligase), *SDHA/SDHB/SDHC* (Succinate dehydrogenase complex), *MDH2* (Malate dehydrogenase), *FH* (Fumarase), *PDHA1/PDHB* (Pyruvate dehydrogenase), *DLD* (Dihydrolipoamide dehydrogenase), *DLAT* (Dihydrolipoamide S-acetyltransferase), DLST (Dihydrolipoamide S-succinyltransferase) and 5) metabolic intermediates that believed to induce epigenetic changes: *FH* gene and *SDHA/SDHB* (blue rectangle).

## 4. Discussion

In this study, we have shown that FFPE core needle biopsy tissues are suitable for spatial transcriptomic analysis, and can uncover the transcriptomic signatures, signaling pathways, and spatially resolved immune landscapes in human kidney allograft rejection. We demonstrated that non-rejection, active AMR, acute TCMR and chronic active AMR have distinct transcriptomic signatures (Fig. 2). We identified distinct subclusters of monocytes and macrophages with high *FCGR3A* expression in C4d-positive active AMR and TCMR, respectively (Fig. 4). The spatial distribution of these distinct clusters corresponded to the characteristic histopathological features of active AMR and acute TCMR, respectively (Fig. 5). Functional pathway and gene network analysis showed upregulation of key pathways that are associated with both metabolic changes in trained immunity and various immune responses, particularly those involving innate immunity (Fig. 6). Moreover, key markers associated with monocytes/macrophage activation and function in both transplant rejection and trained immunity within the innate alloantigen recognition pathway showed significantly increased *CD47* and notably increased *SIPR*α in the C4d-positive active AMR case, while being less prominent in acute TCMR cases (Fig. 7). Finally, our study revealed that the metabolic markers associated with trained innate immunity exhibited distinct expression patterns in groups experiencing rejection compared to those without rejection (Fig. 8). These findings are summarized in Figure 9. To our knowledge, this is the first report of using spatial transcriptomics to evaluate different rejection types of FFPE core needle biopsies from human kidney allografts. Our findings compliment the transcript signatures identified through bulk transcriptome microarrays and scRNA seq, while also providing additional valuable spatial information.

**Figure 9:**
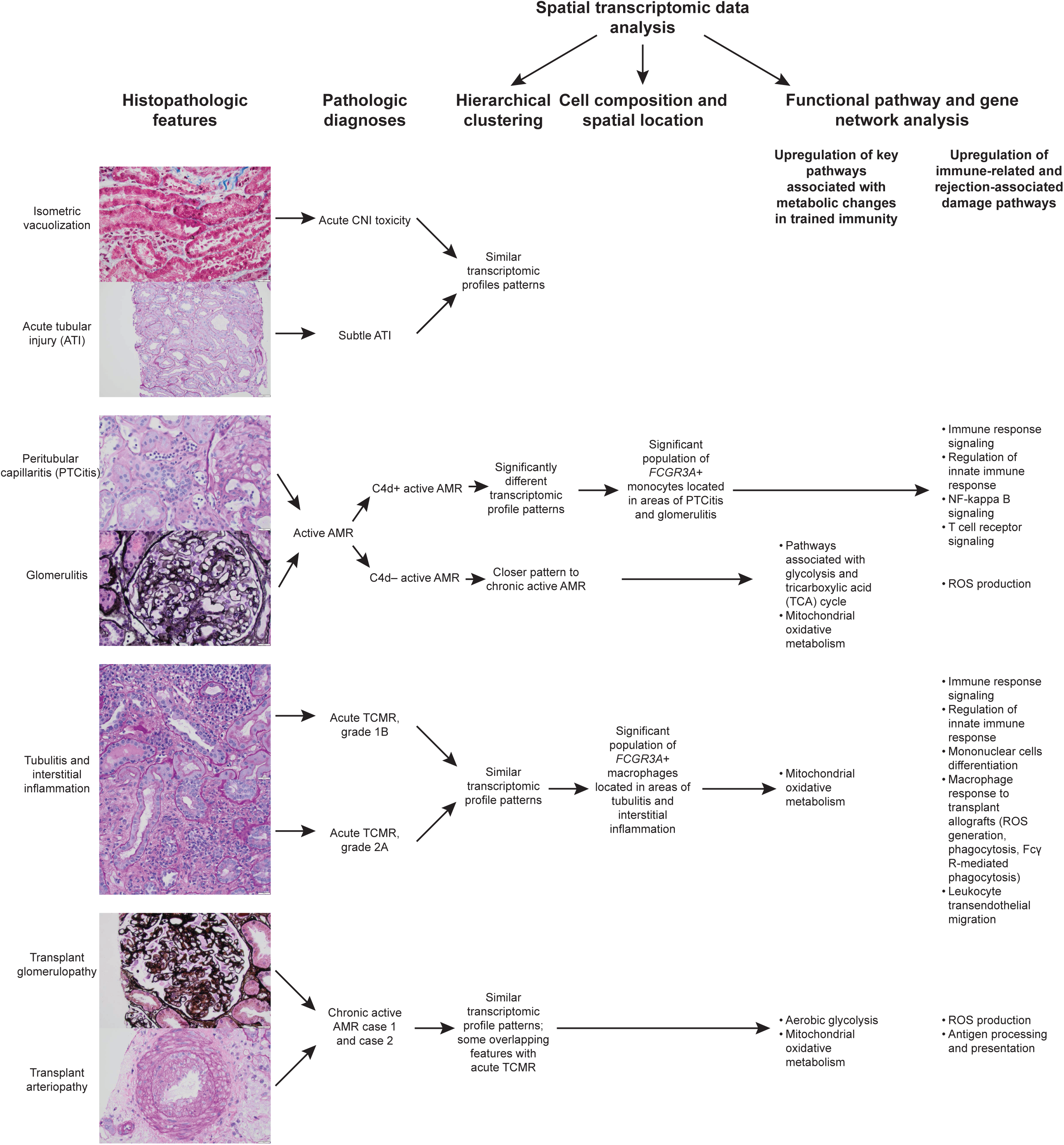
Integration of histopathology, pathological diagnosis, and spatial transcriptomics in eight kidney transplant cases. This figure presents a comprehensive analysis of these kidney transplant cases, illustrating the progression from histopathological features to pathological diagnosis and spatial transcriptomic analysis. The pathological diagnoses include acute CNI toxicity (characterized by isometric vacuolization), active AMR (featuring peritubular capillaritis and glomerulitis, subclassified into C4d-positive and C4d-negative variants), acute TCMR (characterized by tubulitis and interstitial inflammation), and chronic active AMR (featuring transplant glomerulopathy and transplant arteriopathy). The figure demonstrates how these distinct histopathological features inform the pathological diagnosis and how spatial transcriptomics provide additional molecular insights into each condition, offering a multi-layered view of kidney transplant pathology.

Bulk transcriptome microarrays, such as MMDX, have been applied to assist in the clinical diagnosis of rejection. However, these methods typically require relatively large tissue volumes, which are challenging to obtain through core needle biopsies. Moreover, these techniques extract analytes from tissue and sequence them in bulk. Data regarding the type of cells expressing a given transcript, the location of these cells within the tissue, and co-expression of transcripts in the tissue geography are all lost by this bulk preparation. ScRNA seq is a recently developed technology exclusively used in research to analyze gene expression at the individual cell level. While it offers valuable insights into cellular heterogeneity, it has limitations: it typically requires fresh or frozen tissue samples, necessitates a high number of isolated cells that are hard to obtain by core needle biopsy tissue, and loses spatial information. To achieve the required number of isolated cells for scRNA-seq, it will be necessary to pool core needle biopsy tissue samples to perform the experiment. However, this approach will result in the loss of opportunities to study transcriptomic signatures for individual cases. Our approach of using spatial transcriptomics to evaluate rejection on archived FFPE core needle biopsies from human allografts has the potential to bridge the gap between histopathologic and molecular classifications. This approach likely provides more comprehensive information while requiring only minimal tissue input.

Despite advances in immunosuppression regimens used in solid organ transplantation over the past decades, achieving long-term success has been hindered by several challenges, including the need to tailor post-transplant immunosuppression regimens to ensure patient-specific optimization ^39^. Current immunosuppressive treatment regimens only target adaptive immune cells. We are lacking potential biomarkers for innovative immunosuppressive therapies. Although research in the field of innate immunity in transplant immunology has garnered attention in recent years, we lack comprehensive knowledge of the specific transcript signatures associated with innate immune cells during post-transplant events. These events include non-rejection conditions (such as subclinical graft injury, delayed graft function, ATI, CNI toxicity and inflammation below diagnostic thresholds for rejection), early acute rejection, and chronic rejection. Of particular interest are monocytes/macrophages and NK cells, which play critical roles in the innate immune response to transplant allografts by producing proinflammatory factors, killing graft cells, and enhancing the adaptive immune response ^10–14^. Furthermore, organ transplantation induces trained innate immunity, contributing to allograft rejection. However, significant knowledge gaps persist regarding their molecular and cellular mechanism, duration, adaptability and impact on adaptive immunity in human organ transplantation. While clinical trials are ongoing, current immunosuppressive treatment regimens still fail to leverage the potential benefits of modulating the innate immune response. There is a great need to discover potential biomarkers for future innovative immunosuppressive therapies. Our discovery of transcriptomically distinct monocytes/macrophages subclusters and associated signaling pathways in acute rejection, can uncover potential biomarkers, such as *FCGR3A*, for future novel immunosuppressive therapy targets.

An unexpected but potentially important finding was that the C4d-positive active AMR case had significantly different transcriptomic signatures than the C4d-negative active AMR case. Gupta *et al*. found no differences in gene expression between C4d positive and C4d negative biopsies with microvascular inflammation (MVI) >2 using microarrays ^40^. Our findings suggested that spatial transcriptomics may offer a potential advantage over microarray analysis in identifying distinct molecular signatures associated with different morphologic subsets of AMR.

Our study has several limitations. First, the fixed 55-μm diameter map spots on the transcriptomic platform resulted in variable cell densities associated with each barcode. This constraint may have introduced analytical inconsistencies between samples and potentially caused us to overlook less prominent subclusters, such as NK cells. In future experiments, this technical limitation could likely be addressed by applying the newly developed 10x Genomics Visium high definition (HD) spatial transcriptomics technology. Secondly, our study is limited by the number of map spots in capture areas (6 mm x 6 mm). This limitation is due to the nature of kidney needle core biopsy tissue, which is typically small, and the empty gaps between individual tissue cores within the paraffin blocks. To overcome this issue in future studies, we could use larger capture areas (1 cm x 1 cm) and carefully select cases with multiple needle cores. Thirdly, our study was constrained by a small sample size of eight biopsies, due to the high cost of spatial transcriptomics and budgetary limitations. This restricted sample size may account for the lower-than-expected overlap between our FFPE tissue transcriptomic signatures and the previously published MMDX signatures. A larger sample size might have yielded more robust and representative results, potentially increasing the concordance with established findings. Lastly, our ability to investigate complex intercellular communication patterns among monocytes/macrophages and other types of cells, including adaptive immune cells, was limited in this study. This limitation arose because software, such as CellChat, is designed for scRNA-seq data rather than Visium spatial transcriptomics data. In future studies, this constraint could be overcome by integrating scRNA-seq data with spatial transcriptomic analysis, allowing for a more comprehensive examination of cellular interactions within the tissue microenvironment.

In summary, our study demonstrated the feasibility of using archived FFPE core needle biopsy tissues from human kidney allografts for spatial transcriptomic analysis. This will eliminate the need for additional fresh biopsy cores and allows us to study archived specimens with minimal tissue input, enabling the tracking of disease progression at the molecular level. Our discovery of the distinct monocyte/macrophage subclusters in AMR and TCMR at the transcriptomic level may uncover potential cellular targets for developing innovative immunosuppressive therapies.

## Acknowledgement

We appreciate Histology Laboratory and California Tumor Tissue Registry, Department of Pathology and Human Anatomy at Loma Linda University, Loma Linda, California, for their assistance with FFPE tissue processing. We appreciate the support provided by Dr. Paul Herrmann, the Chairman of the Department of Pathology and Human Anatomy. We also thank the Integrative Genomics Core at City of Hope for providing technological and experimental support.

## Data availability statement

The data that supports the findings of this study are available on request from the corresponding author upon reasonable request.

## Abbreviation

PFKFB1: 6-Phosphofructo-2-Kinase/Fructose-2,6-Biphosphatase 1
ACO1/ACO2: Aconitase
AMR: Active antibody mediated rejection
TCMR: Acute cell mediated rejection
ATI: Acute tubular injury
ATP: Adenosine triphosphate
ALDOA: Aldolase A
APC: Antigen-presenting cells
B-HOT: Banff Human Organ Transplant Gene
CNI: Calcineurin inhibitor
CS: Citrate synthase
CLEC7A: C-type lectin domain family 7 member A
DEGs: Differentially expressed genes
DLD: Dihydrolipoamide dehydrogenase
DLAT: Dihydrolipoamide S-acetyltransferase
DLST: Dihydrolipoamide S-succinyltransferase
DV200: Distribution Value 200
DSA: Donor-specific antibodies
ENO1: Enolase 1
FDR: False discovery rate
FCGR3A: Fc gamma receptor IIIA
FFPE: Formalin-fixed paraffin-embedded
FH: Fumarase
GO: Gene Ontology
G6PC: Glucose-6-phosphatase
GPI: Glucose-6-phosphate isomerase
GAPDH: Glyceraldehyde-3-phosphate dehydrogenase
H&E: Hematoxylin and eosin-stained
HK1: Hexokinase 1
HD: High definition
HIF1A: Hypoxia-Inducible Factor 1-alpha
IGF1R: Insulin Like Growth Factor 1 Receptor
IL1R1: Interleukin 1 Receptor Type 1
IDH1/IDH2: Isocitrate dehydrogenase
KEGG: Kyoto Encyclopedia of Genes and Genomes
LDHA: Lactate dehydrogenase A
LILRA: Leukocyte immunoglobulin-like receptor A
MDH2: Malate dehydrogenase
MMDX: Molecular Microscope
MVI: Microvascular inflammation
(NK) cells: Natural killer
NOD2: Nucleotide Binding Oligomerization Domain Containing 2
PPARGC1A: Peroxisome proliferator-activated receptor-γ coactivator 1-α
PGK1: Phosphoglycerate kinase 1
PCA: Principal component analysis
PDHA1/PDHB: Pyruvate dehydrogenase
PKM: Pyruvate kinase
ROS and RNS: Reactive oxygen and nitrogen species
scRNA seq: Single-cell RNA sequencing
SIPRα: Signal-regulatory protein-α
SDHA/SDHB/SDHC: Succinate dehydrogenase complex
SUCLG1/SUCLG2: Succinyl-CoA ligase
TCA: Tricarboxylic acid
t-SNE: t-distributed stochastic neighbor embedding
UMAP: Uniform Manifold Approximation and Projection
YY1: Yin Yang 1
OGDH: α-ketoglutarate dehydrogenase

---

**Supplementary Figure 1: Fc gamma receptor IIIA (*FCGR3A*) expression of each case using Uniform manifold approximation and projection (UMAP) visualization**. *FCGR3A* was significantly upregulated in both C4d-positive active antibody mediated rejection (AMR) and acute cell mediated rejection (TCMR) cases, while non-rejection cases and C4d-negative active AMR exhibited very low expression levels. Chronic active AMR cases showed intermediate expression levels. The color intensity in the visualization represents *FCGR3A* gene expression levels, with red shades indicating higher expression and yellow shades indicating lower expression. CNI: acute calcineurin inhibitors; ATI: toxicity acute tubular injury.

