## Supplementary figures and images for "Spatial Transcriptomics Using Archived Formalin-Fixed Paraffin-Embedded Core Needle Biopsy Tissues Revealed Unique Transcriptomic Signatures in Kidney Transplant Rejections"

### Supplemental Table 1

#### Supplementary Figure 3

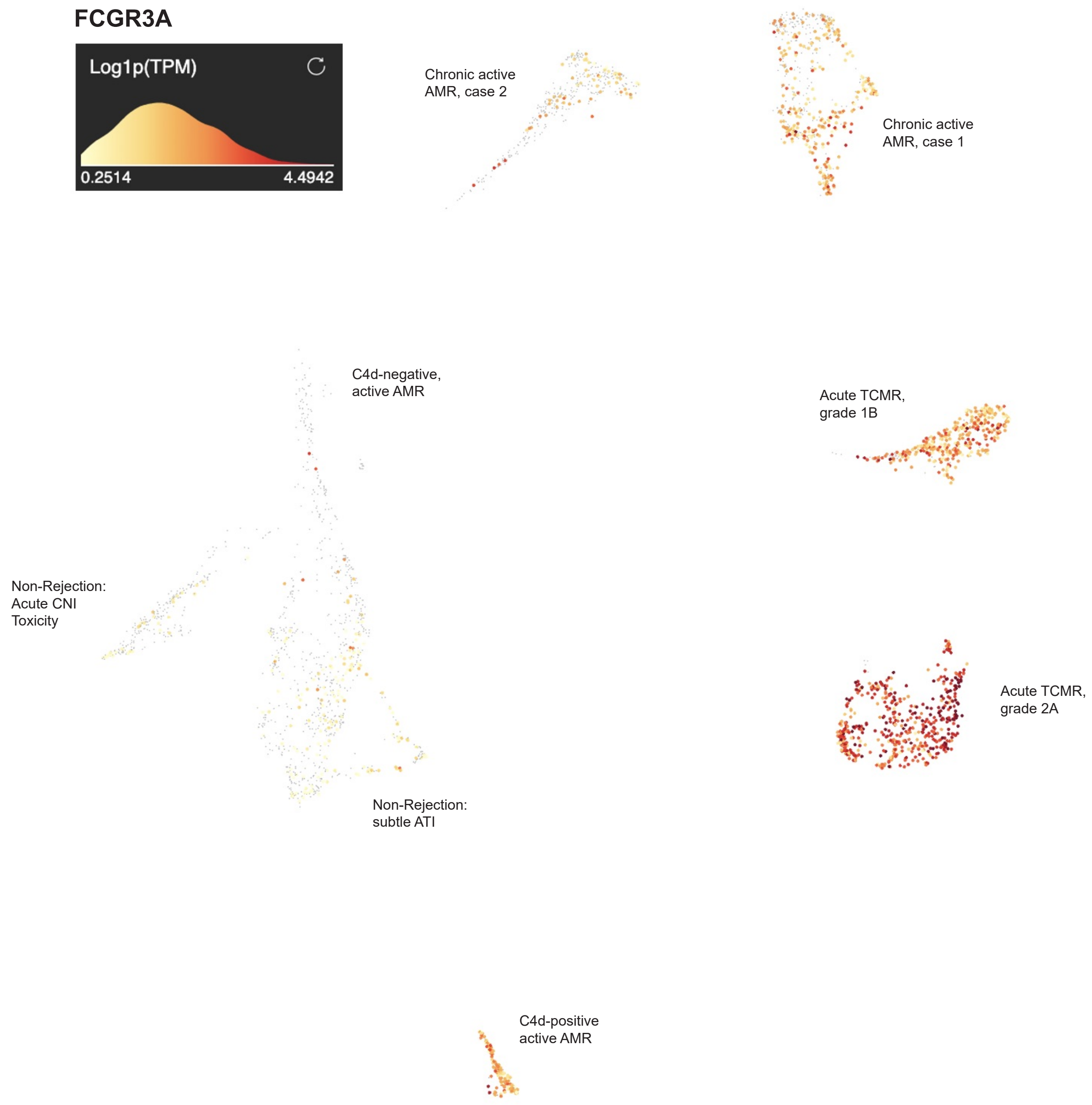
